# Mechanistic Deep Learning Framework on Cell Traits Derived from Single-Cell Mass Cytometry Data

**DOI:** 10.1101/2022.08.22.504669

**Authors:** Bo Wang, Sandra S. Zinkel, Eric R. Gamazon

## Abstract

Germline genetic variations can alter cellular differentiation, potentially impacting the response of immune cells to inflammatory challenges. Current variant- and gene-based methods in mouse and human models have established associations with disease phenotypes; however, the underlying mechanisms at the cellular level are less well-understood. Immunophenotyping by multi-parameter flow cytometry, and more recently mass cytometry, has allowed high-resolution identification and characterization of hematopoietic cells. The obtained characterization yields increased dimensionality; however, conventional analysis workflows have been inefficient, incomplete, or unreliable. In this work, we develop a comprehensive machine learning framework – MDL4Cyto – that is tailored to the analysis of mass cytometry data, incorporating statistical, unsupervised learning, and supervised learning models. The statistical modeling can be used to illuminate cell fate decision and cell-type dynamics. The unsupervised learning models along with complementary marker enrichment analyses highlight genetic perturbations that are significantly associated with alterations in cell populations in the hematopoietic system. Furthermore, our supervised learning models, including deep learning and tree-based algorithms, address the bottleneck to data pre-processing that characterizes conventional workflows and generate inferences (e.g., on marker/cell-type interactions) from raw experimental characterization. Notably, we reveal a close relationship among network design, prediction performance, and the underlying biological context. We show that the network architecture extracted from the differentiation cascade of the investigated biological system yields enhanced prediction performance. The presented methodology will enable new insights into hematopoietic differentiation at baseline and following perturbation.

**Highlights:**

- Analysis pipeline on mass cytometry data with high-performance implementation of statistical, unsupervised learning, and supervised learning models
- Concordance of machine learning results with biological contexts
- Biologically-informed neural network designs enhance prediction performance

**Graphical Abstract:** 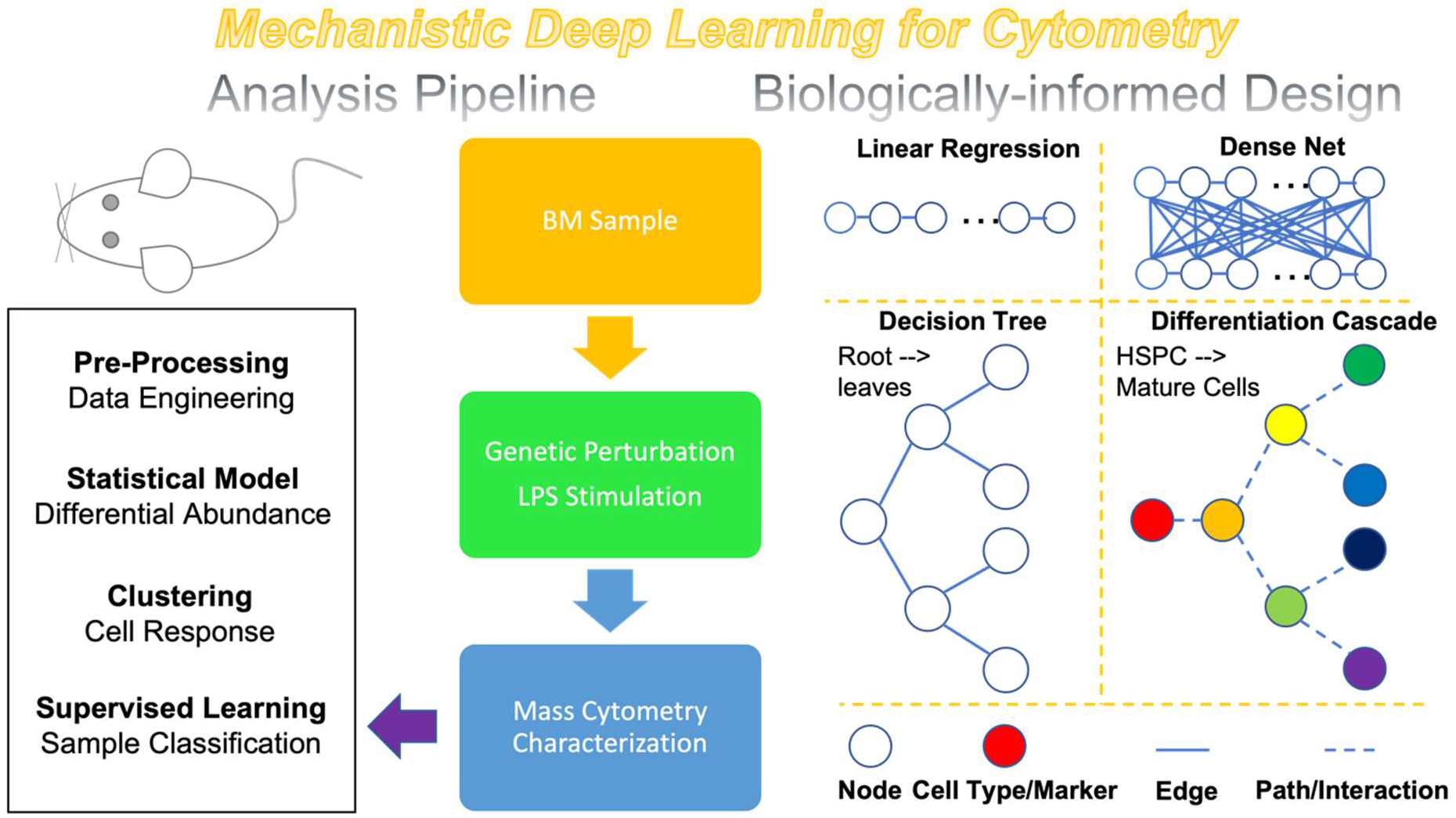

## Introduction

Hematopoiesis is a hierarchical and dynamic process. Proper blood cell development requires highly coordinated and tightly regulated cytokine signals, affecting cell and stage-appropriate gene expression. Genetically modified mouse models are an invaluable model system for studying hematopoiesis, due to their defined genetic background, wealth of reagents to precisely define hematopoietic cell populations, ability to undergo precise genetic modification, and the wealth of data and assays arising from decades of study. Cytometry tools have proved to be essential in simultaneously analyzing labeled molecules at the single-cell level and, consequently, have been widely implemented in basic and clinical research^1^. While traditional fluorescence cytometers can detect up to 35 markers, the technique is limited by spectral overlap of fluorophores. Single-cell mass cytometry (Cytometry by Time-Of-Flight [CyTOF]) enables multiple-parameter tracking by the use of metal-isotope-tagged antibodies, overcoming the inherent limitations of spectral overlap. Therefore, a more comprehensive panel of cellular functions (achieved from significantly expanded feature capacity^2^) along with intracellular signaling states can be explored to identify disease-associated changes or functionally-relevant states; differential expression under external perturbation or intracellular response is more easily elucidated^3^.

Insights from cytometry characterization are traditionally extracted from manual gating. Automated workflows such as FlowCore^4^, PhenoGraph^5^ or the recently reported Tracking Responders EXpanding (T-REX)^6^ have been introduced with significantly enhanced processing efficiency. However, these pipelines are still limited by expert intervention into (manual) data pre-processing, where key thresholds for transformation, gating or cell type clustering are tailored to the investigated systems. Recently, machine learning algorithms have attracted significant attention for their extensive application to the analysis of cytometry data^6–8^. Specifically, beyond functioning as embedded algorithms in pipelines, machine learning tools provide “end-to-end” solutions, generating inferences or predictions directly from raw experimental characterization, with potential to uncover novel associations that are obscured by conventional gating strategies.

Hematopoiesis has served as a paradigm of stem cell biology and cellular differentiation^9^. Genetic perturbations within cells (in particular, Hematopoietic Stem and Progenitor Cells [HSPCs]) induce challenges to cellular functions, thus leading to various diseases or conditions, e.g., Clonal Hematopoiesis of Indeterminate Potential (CHIP), Myelodysplastic Syndromes (MDS), and Acute Myeloid Leukemia (AML). Tet2 inactivating mutations are among the early mutations that initiate clonal expansion and progression to MDS or AML. As the predominant Tet gene in the hematopoietic tissue (especially in HSCs), Tet2 functions as a tumor suppressor whose haplo-insufficiency initiates myeloid and lymphoid transformations^8^. Tet2 has been shown to regulate resolution of inflammatory responses. Tet2 KO mice and cell lines display prolonged and enhanced inflammatory cytokine signaling, which impacts HSPC differentiation.

Hematopoietic homeostasis is further controlled through tight regulation of programmed cell death. The two major types of cell death, apoptosis (programmed cell death) and necrosis (regulated necrotic cell death), maintain homeostasis by downregulating detrimental inflammation and prevent transformation by allowing efficient removal of damaged cells. It has been widely recognized that the genes in the Bcl-2 family are critical in regulating apoptosis^10,11^, comprised of both pro-survival and pro-death members. The functions of BID (pro-apoptotic BH3-only member) were investigated in our recent human phenome-wide association study (PheWAS)^12,13^. In follow-up studies, we evaluated the multidomain effectors Bax, Bak, and Bid in murine models^14^. We found that deletion of Bax/Bak inhibits both bone marrow apoptosis and bone marrow necroptosis. Further deletion of Bid (TKO) leads to robust activation of necroptosis, which results in a phenotype characterized by dysplasia and cytopenia, resembling the bone marrow failure disorder MDS^14^.

To address the limitations of conventional cytometry data analysis pipelines^5–8,15^, we developed a comprehensive *Mechanistic Deep Learning for Cytometry* (MDL4Cyto) framework, implementing statistical, unsupervised learning, and supervised learning models. The framework was evaluated against bone marrow samples of murine models that have been genetically perturbed in the hematopoietic system, including Wild Type (WT), *VavCreBaxBakBid* triple-knockout (TKO), and *VavCreTet2* knockout (Tet2 KO) mice, and validated with biological observations. Notably, building on our previous work^16,17^, we designed the architectures of the neural networks using the intrinsic hierarchy of the hematopoietic system as a guide, establishing a *biological feature – network architecture – phenotype* relationship. We show that biologically-informed network designs could function as probes on the regulatory cascade with prediction performance as the evaluation metric. Our framework may inform future machine learning implementations to address fundamental questions in cell biology.

## Results

### Overview of mass cytometry data and MDL4Cyto framework

Mice from three genetic murine models – WT, Tet2 and TKO – were treated with Phosphate-buffered saline (PBS) or lipopolysaccharide (LPS). LPS is a major surface membrane polysaccharide found on Gram-negative bacteria and is a typical pathogen-associated molecular pattern (PAMP). Inflammation triggered by LPS stimulation activates a robust innate immune response and can induce the proliferation of hematopoietic progenitor cells^18^ and the production of cytokines such as TNFα, interleukin and IL-2. Bone marrow from these mice was subsequently harvested and processed for cytometry experiments. Besides genetic perturbations, all the samples challenged by LPS stimulation were compared to the PBS vehicle control (Figure 1A). Across different types of samples (WT-PBS, WT-LPS, Tet2-PBS, Tet2-LPS, TKO-PBS, and TKO-LPS), lineage marker groups (stem cells, pan leukocytes, T lymphocytes, mature myeloid cells, B lymphocytes, and erythrocytes) and functional marker groups (cytokines, cell death, and proliferation), were tagged and monitored (Table S1). As sample data profiles obtained from the mass cytometry measurements varied in length, we imputed data points at each of the six sample data profiles via binning methods with 30 seconds.

**Figure 1.**
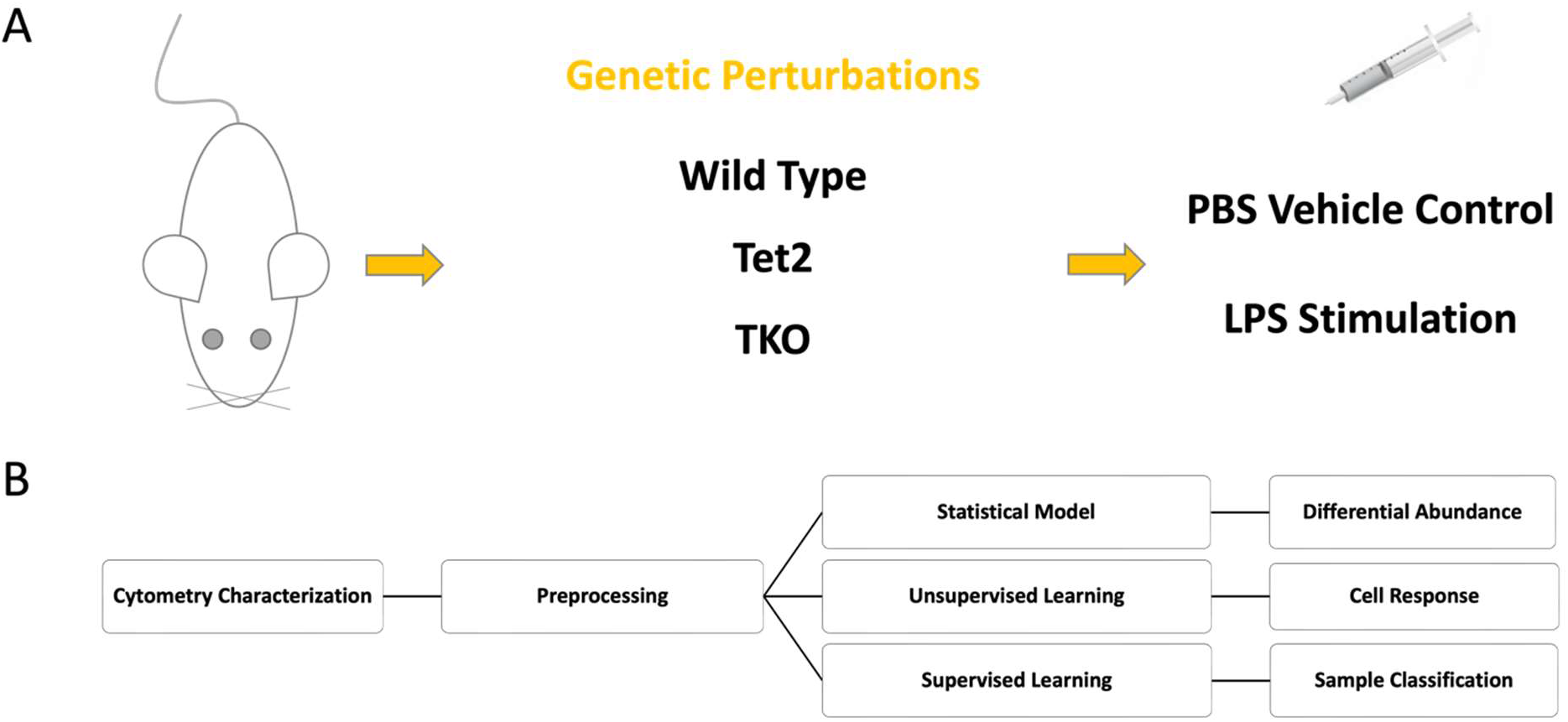
MDL4Cyto pipeline for analysis of single-cell mass cytometry data. Bone marrow samples (**A**) were harvested from genetically perturbed murine models (Wild Type [WT], *VavCreBaxBakBid* triple-knockout [TKO], and *VavCreTet2* knockout [Tet2 KO]), which had been treated with LPS stimulation or PBS vehicle control. Statistical, unsupervised, and supervised learning models were implemented in our *Mechanistic Deep Learning for Cytometry* (MDL4Cyto) framework. The analysis pipeline **(B)** was benchmarked, using the reference dataset obtained from mass cytometry characterization, for the tasks of differential abundance analysis, cell response detection, and sample type classification.

We developed MDL4Cyto, an analysis pipeline for mass cytometry data (Figure 1B and **Methods**), implementing statistical, unsupervised, and supervised learning models. Statistical tests on the cell count of key lineage and functional markers were performed, aiming for a glimpse of the underlying biology. As both genetic perturbations and LPS stimulation may induce inflammatory changes in hematopoiesis, we applied Generalized Mixed-Effects Model (GMM) estimation to determine the extent to which cell count was statistically altered by these modifications. Two established workflows, namely PhenoGraph^5^ and T-REX^6^, were benchmarked as established pipelines capable of annotating cell populations or capturing cell responses. In addition, supervised learning models were explicitly designed to probe marker/cell-type interactions within the regulatory cascade, as well as to predict sample type directly from the raw mass cytometry characterization. Nine machine learning models were evaluated, including Linear Regression (LR), Dense Neural Network (DNN), Boosted Tree (BT), Random Forest (RF), stacked Dense Neural Network - Random Forest (DNN-RF), Convolutional Neural Network in One Dimension (CNN-1D), Convolutional Neural Network in Two Dimensions (CNN-2D), Recurrent Neural Network with Long Short-Term Memory cell (RNN-LSTM), and Recurrent Neural Network with Gated Recurrent Units cell (RNN-GRU).

### Statistical modeling of genetically-determined or condition-dependent effects

The cell count summary statistic, which was calculated on the reference dataset, was found to be altered by genetic perturbations and/or LPS stimulation (Figure S1). Clear differences were observed for all markers. As reported in our previous work^14^ and other studies^19^, the TKO sample leads to necrosis with increasing production of cytokines. Enhanced inflammatory signaling was also observed in Tet2-deficient samples, as has also been previously described^19^. We observed a statistically significant difference in cell count for Sca1 (LSK cells), CD45 (leukocytes), TNFα (cytokines), and Caspase 3 (cell death indicator) with respect to genetic perturbation (Figure 2A). Markers of cytokines were found to be highly correlated with other surface markers, especially under LPS stimulation (Figure 2B).

**Figure 2.**
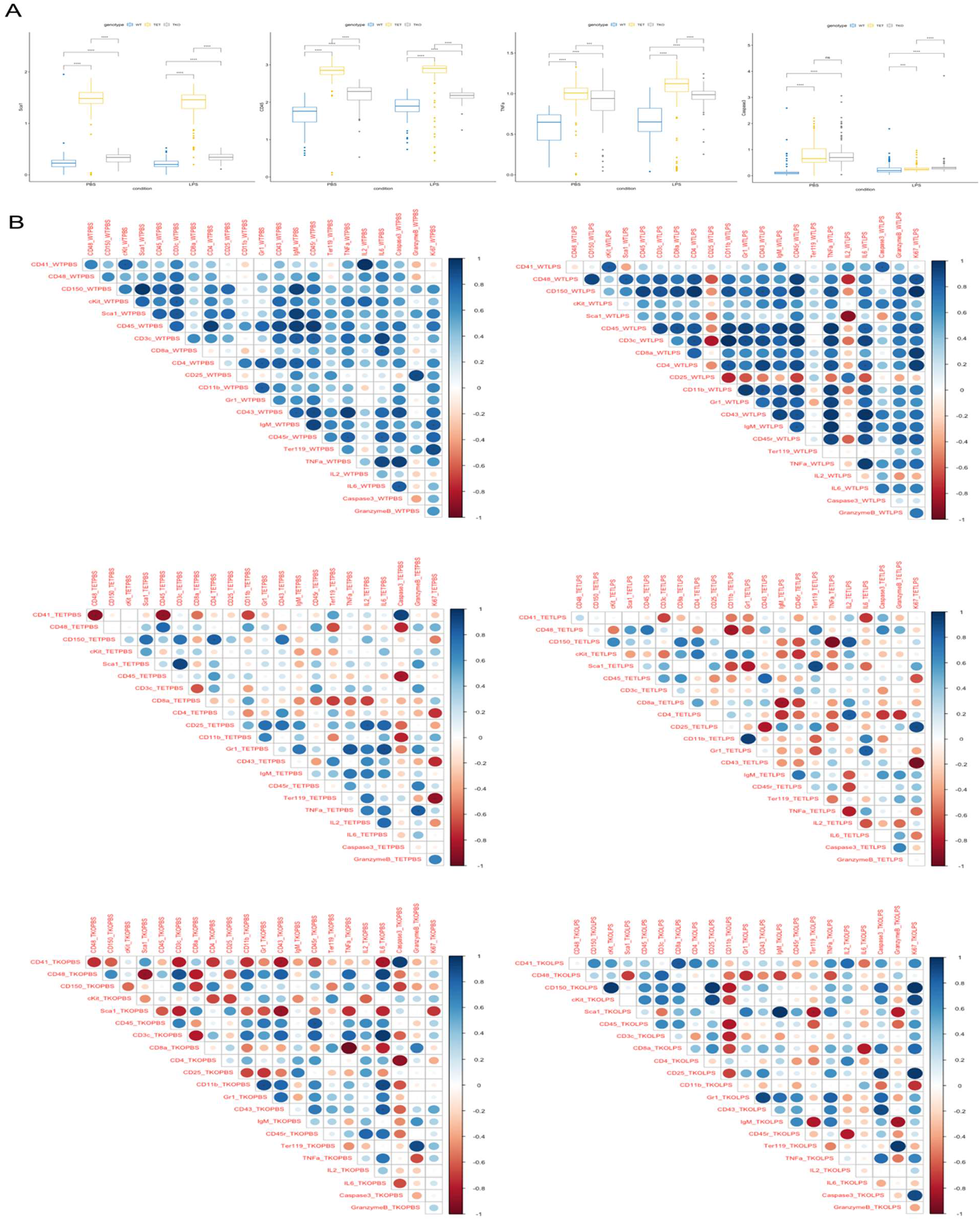
Impact of genetic perturbations on cell populations and correlations among the markers. A statistically significant change in cell count was observed for Sca1 (LSK cells), CD45 (leukocytes), TNFα (cytokines), and Caspase 3 (cell death indicator) in both the PBS vehicle control and LPS stimulation condition, demonstrating genetic perturbations altering cell populations **(A)**. Cytokines from genetic knockout samples were highly correlated with other surface markers, in particular under LPS stimulation **(B)**, which is consistent with the biological context. (Wilcoxon test was performed for the pairwise comparisons; Pearson correlation was calculated for the markers along with 95% confidence interval. *p<.05, **p<0.01, ***p<0.001, ****p<0.0001).

The data profiles obtained from cytometry characterizations are dynamical panels, i.e., have a time component. To estimate the overall effects of genotype and condition, we analyzed the progression of marker Intensity for Ki67 from the aggregated data panel with 1 hour binning window (Figure 3 and **Methods**). Ki67 is a proliferation protein that is increased in the S phase of the cell cycle and indicates proliferating cells. We observed progression divergence according to genotype (Figure 3A) and differentiated marker intensity between conditions (PBS vs LPS, Figure 3B). This is consistent with our understanding that genetic perturbations and LPS stimulation impose pressures on differentiation and cell cycle as dynamical processes. To achieve a better quantitative estimate of the contributions arising from genotype and condition, we implemented GMM estimation. Besides the fixed and/or interaction terms (**Methods**), the time term was considered in the model (Figure 3C). We applied corrected Akaike’s Information Criterion (AICc) and Bayesian Information Criterion (BIC) for model evaluation and Marginal R squared (R2m) for variance coverage. Models built from genotype demonstrated the best performance, which corroborates the fact that genetic perturbations have significant impact on cellular differentiation and function. We note that the statistical modeling implemented in MDL4Cyto (Figure 1B and **Methods**) is broadly applicable, which can facilitate the study of cell fate decision and cell-type dynamics.

**Figure 3.**
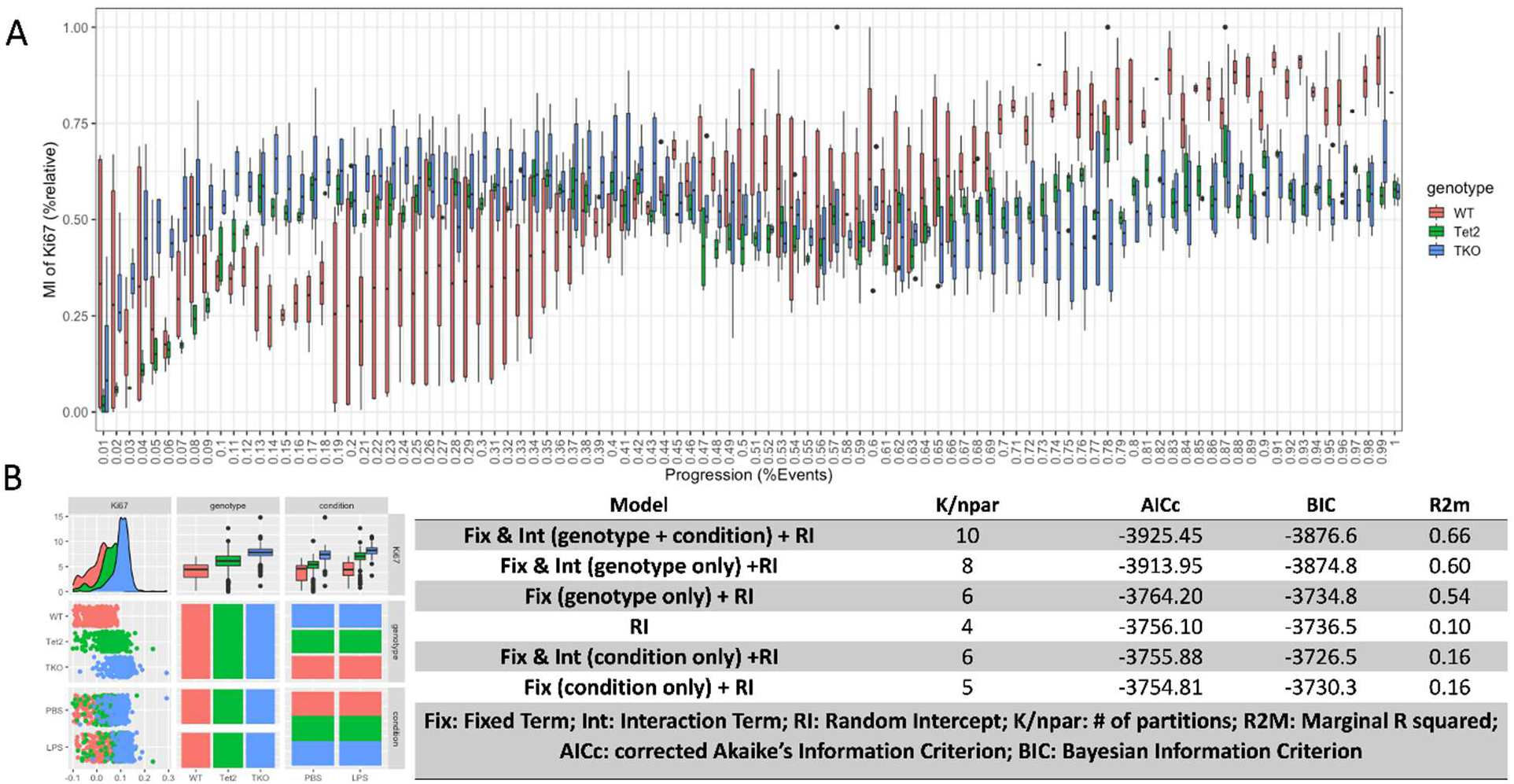
Analysis of dynamic data panel. Progression (%Events over the course of time) **(A)** and histogram **(B)** of cell count of proliferation marker Ki67 diverged with genetic background and shifted with condition. We implemented Generalized Mixed-Effects Model (GMM) estimation that considered the time term in addition to a fixed term and interaction term. While using genotype and condition as main parameters, several models were constructed, as represented by the equations in **(C)**. The performance of each model was further evaluated using the corrected Akaike’s Information Criterion (AICc), Bayesian Information Criterion (BIC), and Marginal R squared (R2m). Genotype was identified as an important feature.

### Quantifying cell response and population alteration via unsupervised learning

Identifying cell response from cell populations has been a key interest in mass cytometry analysis. Hematopoietic responses, induced either by genetic perturbations or by pathogen-like stimulation (LPS), were analyzed via pairwise comparisons, including WT-PBS vs TKO-PBS, WT-LPS vs TKO-LPS, WT-PBS vs Tet2-PBS, WT-LPS vs Tet2-LPS. Two established unsupervised learning algorithms, T-REX and PhenoGraph, were benchmarked in clustering our reference dataset (see Table S3 for the hyperparameters). Both algorithms apply k-nearest neighbors (kNN) for clustering and Uniform Manifold Approximation and Projection (UMAP) for dimension reduction and visualization. As illustrated in the UMAP plot, genetic perturbations induced alterations in cell populations (Figure 4A&4B). Notably, differential cell count of markers under a genetic perturbation, including Sca1, CD45, TNFα, and Caspase 3, could be seen in the mapping of cell count to the clusters (Figure S2, T-REX algorithm). These results are consistent with the comparisons from our statistical tests (Figure 2A). Compared to conventional methods, unsupervised learning showed a unique advantage in efficiently capturing changes in cell populations. In our framework, cell subpopulations within the clusters were further investigated in two ways. While reference datasets trained here contained only a single sample for each type, we created two pseudo-replicates from a single sample (**Methods**) for the PhenoGraph algorithm, thus enabling differential abundance analysis for the markers within each cluster (Figure 4C). In addition, cell populations from the T-REX algorithm were screened and enriched through Density-Based Spatial Clustering and Application with Noise (DBSCAN) and Marker Enrichment Modeling (MEM)^20^ (Figure 4D). Strikingly, in contrast to the standard T-REX method, our framework uncovered a strong signal from the cytokines, i.e., IL6 and TNFα, from the differential abundance analysis (Figure 4C). This observation could well be attributed to the known LPS-driven inflammation, as demonstrated by the enrichment analysis from MEM (Figure 4D). Collectively, the TNFα and IL6 differential abundance results are consistent with myeloid-driven or innate inflammation.

**Figure 4.**
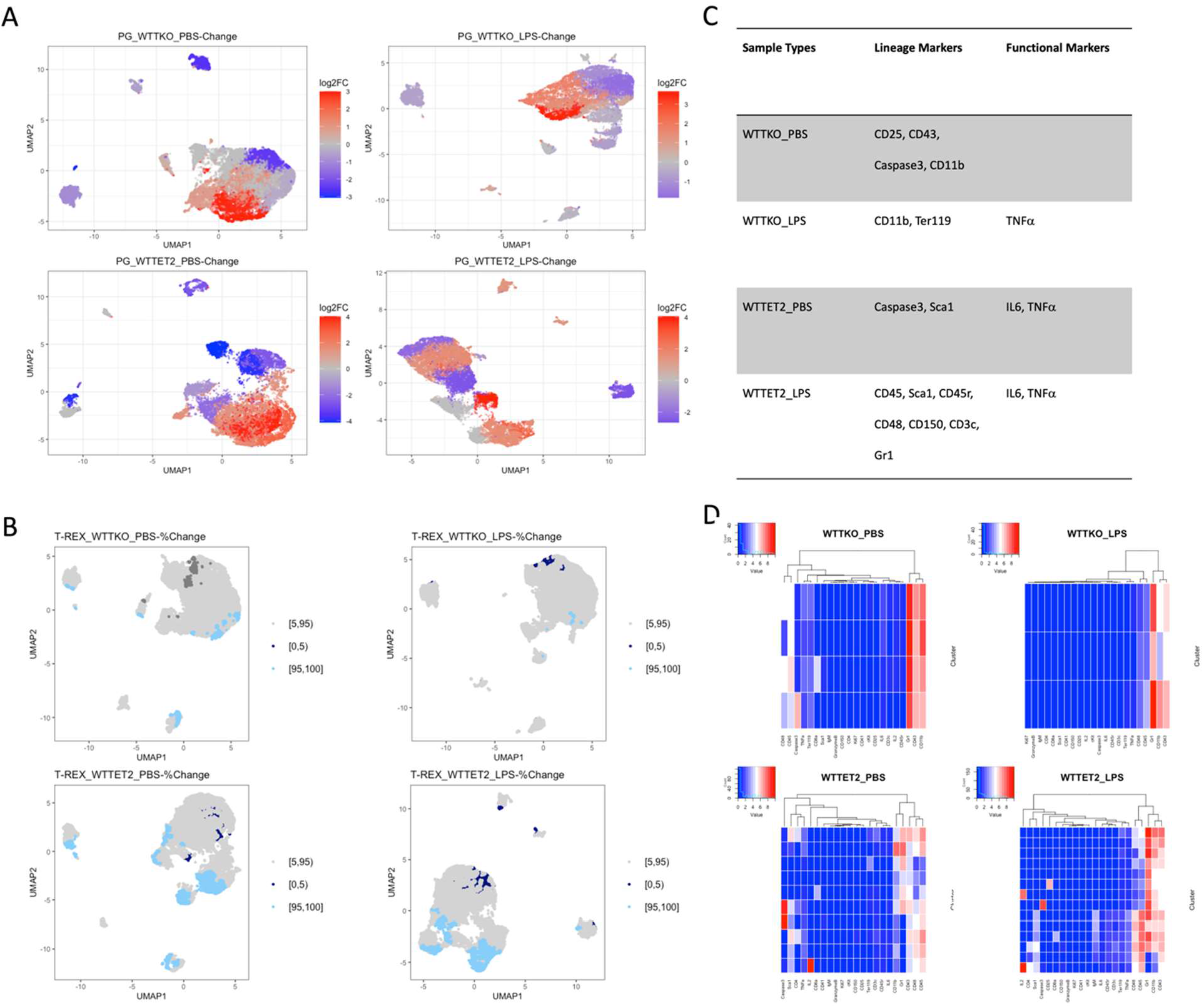
Cell response identified by the unsupervised learning. Clustering from Phenograph **(A)** and T-REX **(B)** algorithms was performed on the paired genetically perturbed samples. PBS vehicle control and LPS stimulation were separately explored and compared. Using both algorithms, we found that the genetic perturbations resulted in altered cell populations. In our framework, clusters of significant fold changes identified from PhenoGraph were further explored for differential abundance of markers. Markers with significant responses that were captured in more than two clusters were identified **(C)**. In addition, cell populations obtained from T-REX were screened for those with significant changes and enriched through DBSCAN and MEM **(D)**.

### Probing the regulatory cascade with biologically-informed neural networks

The hematopoietic system has classically been represented as a rigid hierarchical tree, illustrating HSC progression in stages, from multipotent to differentiated cell types. Evaluating a “dynamic” model with complex interactions, we interrogated the marker/cell-type interactions within the regulatory cascade from supervised learning models, considering both the network architecture and prediction performance as probes. LPS stimulation impacts all stages of HSC differentiation, thus having a global impact on all surface markers^18^. Therefore, we assigned PBS vehicle control and LPS stimulation as predicted labels to the samples (**Methods**). Twenty-two lineage/functional makers from nine different cell types (Table S1) were encoded as essential numerical features, while genotype was loaded as an external categorical/global feature.

Considering each node as a marker/cell type, their interactions could be represented as network connections. Specifically, a linear regression model reflects the additive effect from markers/cell types; a dense neural network models the complex interactions among markers/cell types; and a tree model considers the binary relations between levels within the hierarchy (Figure 5A). LPS stimulation on the cells (the prediction label) could be treated as a perturbation on the neural network. In this model, HSPCs will be differentiated to the common lymphoid progenitors, unipotent precursor cells as well as megakaryocyte-erythroid progenitor cells, eventually producing mature cells. Informed by the differentiation hierarchy, we implemented Boosted Tree (BT) and Random Forest (RF) models along with two reference models, Linear Regression (LR) and Deep Neural Network (DNN). The four models varied in performance: Linear Regression (LR) achieved a prediction accuracy of 64.8%; the DNN model with three-hidden layers enhanced the accuracy to 68.1%; the Boosted Tree (BT) model further increased the prediction performance to 72.7%; and the Random Forest (RF) models topped the accuracy at 77.13% (Figure 5B). This overall pattern of prediction accuracy (LR < DNN < BT < RF) suggests a non-linear, hierarchical relationship among the features, consistent with the classical hematopoietic hierarchical tree. Interestingly, when we stacked a Deep Neural Network on top of the Random Forest model, where DNN functioned as a denoising head^21^, the new stacked model showed degraded performance in comparison to the RF model alone.

**Figure 5.**
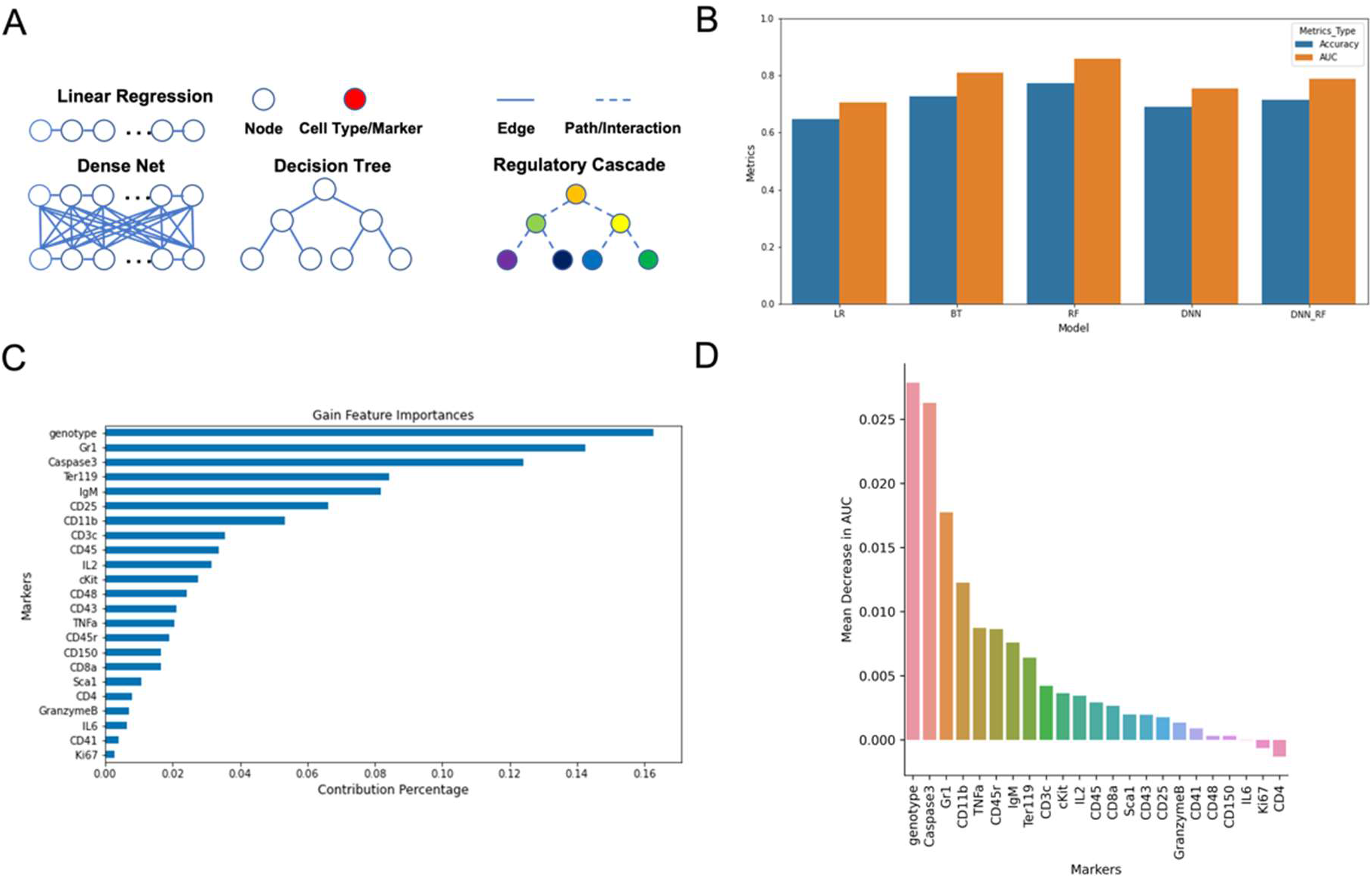
Mechanistic insights extracted from biologically-informed neural networks. Our neural network designs were guided by the biological context, in which we considered each node as a marker. The marker/cell-type interactions could therefore be represented as a network **(A)**, e.g., a decision tree or neural network could be linked to the regulatory cascade. With PBS vehicle control and LPS stimulation as labels, tree-based models (Boosted Tree [BT] and Random Forest [RF]) showed improved performance compared to Linear Regression (LR) and Deep Neural Network (DNN) models as well as DNN-RF stacked models **(B)**, consistent with the hematopoietic tree hierarchy (i.e., representing a sequential binary branching decision process) among the markers. Feature importance of individual markers in the BT model, which implements gain-based feature importance **(C)**, and RF model, which implements mean decrease in AUC **(D)**, were examined. The group of markers with significant contributions would imply an overall cell response under inflammatory (LPS), e.g., pathogen-induced, challenges.

We further considered the feature contributions of individual markers using gain-based feature importance (Figure 5C) and mean decrease in AUC (Figure 5D). Gain-based feature importance reflects the relative contribution of a given feature to the model. Mean decrease in AUC indicates the decline in performance once the corresponding feature has been removed from the training dataset. We hypothesized that markers/features with the top feature contributions are biologically meaningful.

Indeed, the single most important contribution came from the strongest perturbation – genotype. Individual markers, including Caspase3, Gr1, IgM, CD11b, and CD25, that contribute substantially to the prediction of LPS stimulation, are highly biologically relevant in the full differentiation cycle of hematopoiesis. Caspase 3 is an important player in the regulated cell death. Gr1 and CD11b reflect a response from myeloid cells. And Ter119 indicates a strong immune response from the erythrocytes under inflammatory challenges.

### Applying machine learning in sample classification

While conventional methods/pipelines are limited by domain knowledge or manual intervention, a predictive model that bypasses cell clustering or manual gating may provide an end-to-end approach to the analysis of mass cytometry data. Here, we benchmarked a group of supervised learning models, including CNN-1D, CNN-2D, RNN-LSTM, and RNN-GRU, that directly handle the raw cytometry profiles for sample classification, i.e., prediction of clinical or biological outcome (Figure 6A). We established multi-class output labels corresponding to six different sample types, including WT-PBS, WT-LPS, Tet2-PBS, Tet2-LPS, TKO-PBS, and TKO-LPS. We used the curated ensemble of (1 hour) “segments” from the initial bulk samples in evaluating the model performance. We found that CNN and RNN-derived models demonstrated good prediction performance with at least 95% accuracy (Figure 6B). We argue that convolutional layers and recurrent cells are efficient in learning the relevant features. Algorithms such as CNN and LSTM as applied here, with cell count as the trait, captured with greater fidelity the landscape of marker/cell-type interactions than the tree-based discrete models evaluated above.

**Figure 6.**
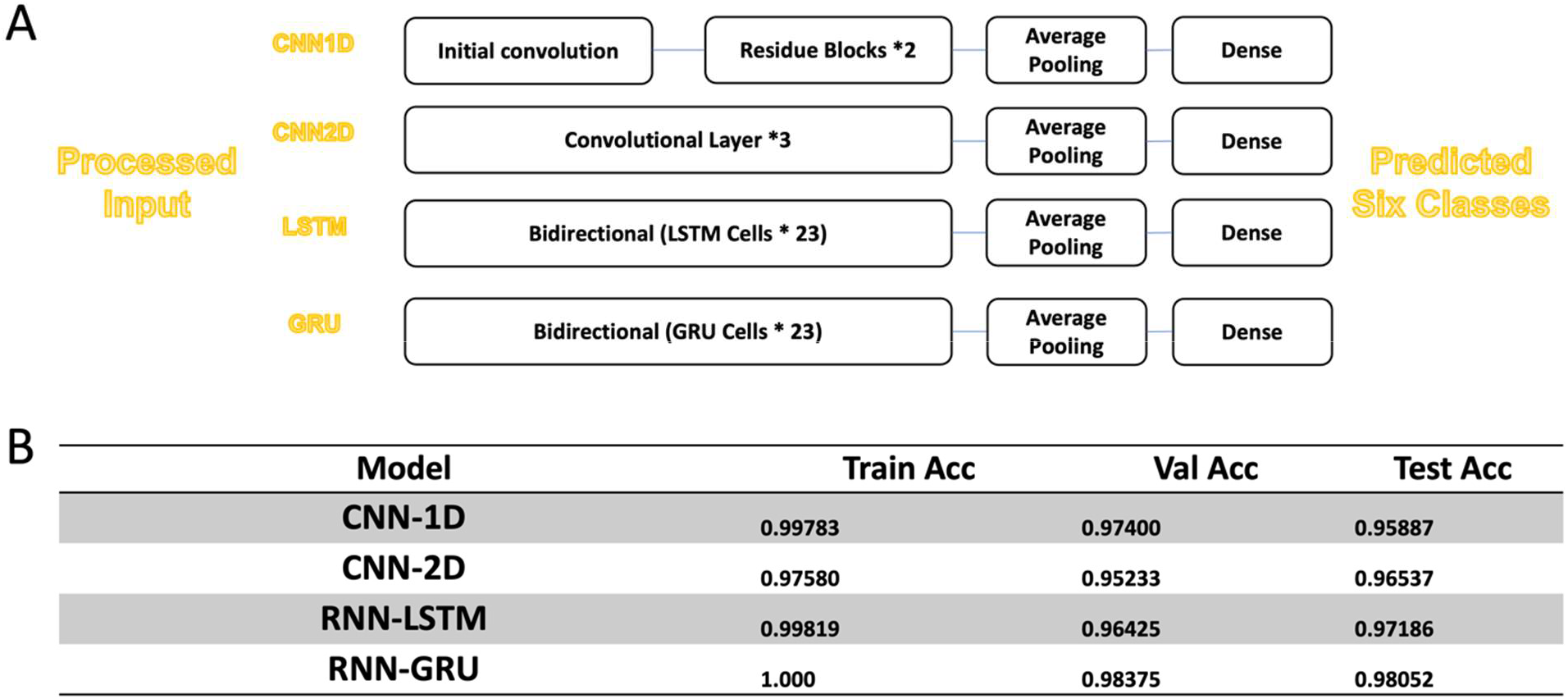
Supervised learning identified sample type with high prediction performance. Convolutional Neural Network (CNN) and Recurrent Neural Network (RNN) derived models (CNN-1D, CNN-2D, RNN-LSTM, RNN-GRU) were benchmarked for the task of sample type classification. The specifics of the architectures are shown in **(A)**. All tested models achieved high performance with over 95% in accuracy **(B)**, demonstrating the importance of learning the complex interactions and high-order features.

## Discussion

The current single-cell genomic revolution has enabled entire systems, such as the hematopoietic system, to be interrogated comprehensively at an unprecedented resolution, revealing a previously unappreciated degree of functional and phenotypic variation within cell compartments. In this work, bone marrow samples were harvested from mice and further characterized in mass cytometry – the baseline murine model (WT) was compared to genetic perturbations, including *Tet2*-knockout (*VavCreTet2*) and *VavCreBaxBakBid* triple-knockout (TKO). LPS served as an external stimulus or an innate immune response, simulating pathogen infection. Statistical modeling and unsupervised learning revealed the direct effect of genetic perturbations. Markers of interest that are typically processed individually could be combined in unsupervised learning models. Importantly, our study highlights a close connection among the intrinsic hierarchy of HSPC differentiation, neural network architecture, and prediction performance. Specifically, Boosted Tree (BT) and Random Forest (RF), which closely resemble the stepwise binary branching decisions of the classical hierarchical model of hematopoiesis, showed superior performance in comparison with Linear Regression (LR), Deep Neural Network (DNN) or even the DNN-RF stacked model. The prediction performance was substantially enhanced relative to neural network models, as the model reflected the cell differentiation cascade. However, as recent studies with functional assays, non-invasive lineage tracing, and omics studies at single-cell resolution have shown^22^, hematopoiesis is a continuous process in which obvious boundaries are lacking between cell populations located at different levels of the hierarchy. Consequently, given their architecture, tree-based models may have an inherent limitation in capturing transition states, even with extensive optimization, constraining their ability to further improve prediction performance. From this perspective, the complex relationships among the cell populations (including feed-back loops in pathways) may be better elucidated by neural networks.

Nevertheless, results of the analyses from the machine learning models were consistent with biological insights such as myeloid-driven or innate inflammation (responses from TNFα and IL6). In summary, the presented framework provides a unifying methodological framework for the analysis of cytometry data, combining data-driven and model-driven approaches. We constructed interpretable models that are anchored in biological context, with implications for ongoing and future studies in single-cell biology.

## Limitations of the study

Machine learning models, including graphical representation learning and tree-based algorithms, were trained in datasets obtained from mass cytometry (CyTOF) characterization. While we designed the neural network architectures with biological information as guide, assumptions of cell types as nodes and edges as their interactions within a graph-based framework are based on the classic discrete cell differentiation model. Cell differentiation is a continuous process, with intermediate states among various levels leading to much more complex networks.

## Methods

### Mass cytometry and murine samples

Three generic murine models were included in this study, namely, wild type (WT), Tet2 knockout (Tet2)^23^, and *VavCreBaxBakBid* triple-knockout (TKO)^14^. Prior to bone marrow isolation, the mice had been treated intraperitoneally with 1.5 mg/kg LPS (L4391; Millipore-Sigma) or PBS for 8 hours. The obtained samples for the six genotype-condition types were subsequently characterized using mass cytometry, according to previously established protocols^24^. Mass cytometry combines inductively coupled plasma time-of-flight mass spectrometry and cytometry. Specifically, mass cytometry staining was performed on whole bone marrow with antibodies conjugated to heavy metals. Cell responses from 9 different cell type and functional categories (stem cells, pan leukocytes, T lymphocytes, mature myeloid cells, B lymphocytes, erythrocytes, cytokines, cell death, and proliferation) were recapitulated by 16 surface/lineage markers and 6 function/intracellular markers. Markers from pre-gating channels (DNA-1 and DNA-2) and instrument channels (Time/Event) were also considered (Table S1). The CyTOF data generated from the six samples consisted of a maximum of 293,778 cells and a minimum of 121,124 cells.

### Data pre-processing

Initial cell count data profiles obtained from the mass cytometry characterization were filtered out for debris, doublets, and dead cells, thus reducing noise. They are represented as dynamical panels on which the *arcsinh* transformation with co-factor = 5 and a binning window of 30 seconds was applied. We sought to analyze the data in a way that maximizes robustness. The processed profiles were sliced into ensembles of “segments” to achieve an optimized tradeoff between computational efficiency and prediction accuracy. We evaluated an array of sizes ([2, 10, 30, 60, 120, 240, 480]), corresponding to 2*30S=1M, 10 * 30S = 5M, 30 * 30S = 15M, 120 * 30S = 1H, 240 * 30S = 2H, & 480 * 30S = 4H sampling intervals (M: Minute, H: Hour), respectively. The obtained pile of “segments” was completely shuffled and re-assembled into new ensembles. We implemented an RNN-LSTM model for this special “hyperparameter search” (see prediction accuracy in Table S2) to determine the optimal segment size, which was 1H. Signals of select markers, including Sca1, CD45, TNFα, Caspase 3, and Ki67, were aggregated and averaged into 1 hour binning window for the statistical test and GMM estimation.

### Mechanistic Deep Learning Framework

#### Statistical test and model

Wilcoxon test was performed on pairwise comparisons of marker intensity. Pearson correlation coefficient was calculated for inter-marker association. We implemented a GMM on the marker Ki67. We considered fixed effects and interactions among time, genotype, and condition (PBS control or LPS stimulation). After testing the random slope of a sample, (1 + Time | Sample), we considered only the random intercept from a sample, i.e., (1 | Sample). The models were evaluated using corrected Akaike’s Information Criterion (AICc), Bayesian Information Criterion (BIC), and Marginal R squared (R2M). The optimal model was found to be given by the following expression:

Marker Intensity = Time + Genotype + Condition + Genotype: Time + Condition: Time + (1 | Sample) + ε

#### Unsupervised learning models

We analyzed two established unsupervised learning pipelines, namely Phenograph^5^ and T-REX^6^, which were applied to the preprocessed CyTOF dataset with binning window 30S. For both pipelines, Uniform Manifold Approximation and Projection (UMAP)^25^ was used for dimensionality reduction; k-nearest neighbors (kNN)^26^ for clustering; Marker Enrichment Modeling (MEM) for generating the enriched heatmap. The applied hyper-parameters are presented in Table S3. For comparison, we screened and enriched the cell populations derived from the T-REX algorithm through Density-Based Spatial Clustering and Application with Noise (DBSCAN) and MEM.

#### Supervised learning models

All the machine learning models were trained using the *TensorFlow (TF)* ecosystem^27^, which we applied to the preprocessed CyTOF dataset (with binning window size of 30S) to perform model training and evaluation. Linear Regression (LR), Deep Neural Network (DNN), Boosted Tree (BT), and Random Forest (RF) models were implemented on the dataset with train and test split of 8:2. The DNN was implemented as a multilayer perceptron. A stacked model of Dense Neural Network and Random Forest (DNN-RF) was constructed following the same splitting. Models implementing a Convolutional Neural Network (CNN) or a Recurrent Neural Network (RNN) in the framework include CNN-1D, CNN-2D, RNN-LSTM, and RNN-GRU. For these four architectures, pre-processed datasets were divided into train, valid, and test data according to 6:2:2 ratio.

We trained the neural networks using mean squared error (as the loss function), *Adam* (an efficient stochastic optimization method), and Leaky ReLU (as the activation function), building on our previous work^16,17^. To reduce the possibility of overfitting, we implemented *dropout* assuming a dropout rate of 20%.

## Acknowledgments

This research is supported by National Institutes of Health (NIH) grants NHGRI R35HG010718, NHGRI R01HG011138, NIA AG068026, and NHLBI R01HL133559.

## Author contributions

B.W., S.S.Z., and E.R.G. conceived the project and designed the study. B.W. performed the analyses and drafted the article. B.W., S.S.Z., and E.R.G. revised the article. S.S.Z. and E.R.G. supervised the project and acquired funding for the study.

## Supplementary Information

**Figure S1.**
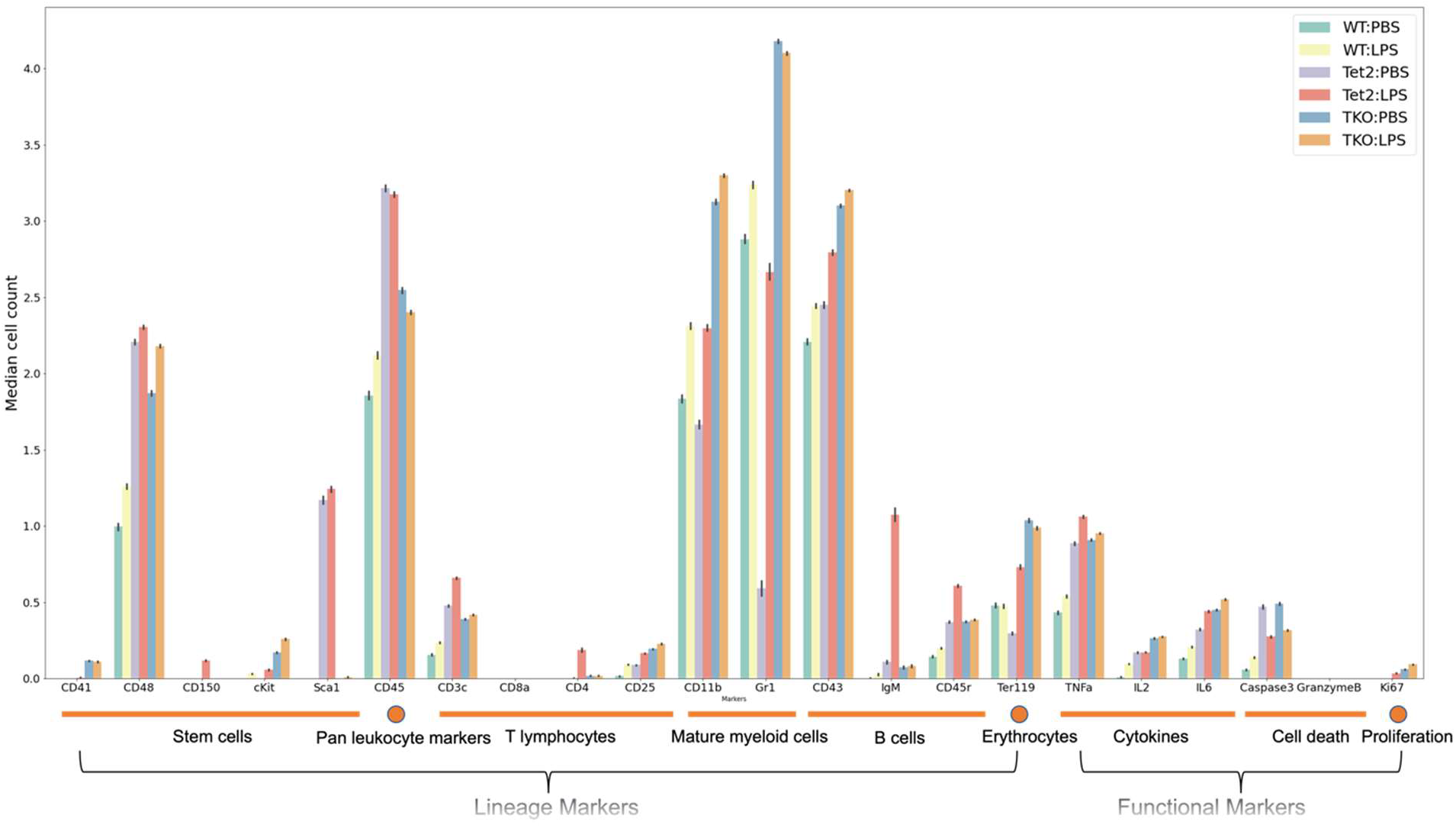
Cell count summary statistics. The sample data profiles (WT-PBS, WT-LPS, Tet2-PBS, Tet2-LPS, TKO-PBS, and TKO-LPS) were obtained from mass cytometry characterization, from which cell count summary statistics were computed. Lineage and functional marker groupings are defined by the cell types.

**Figure S2.**
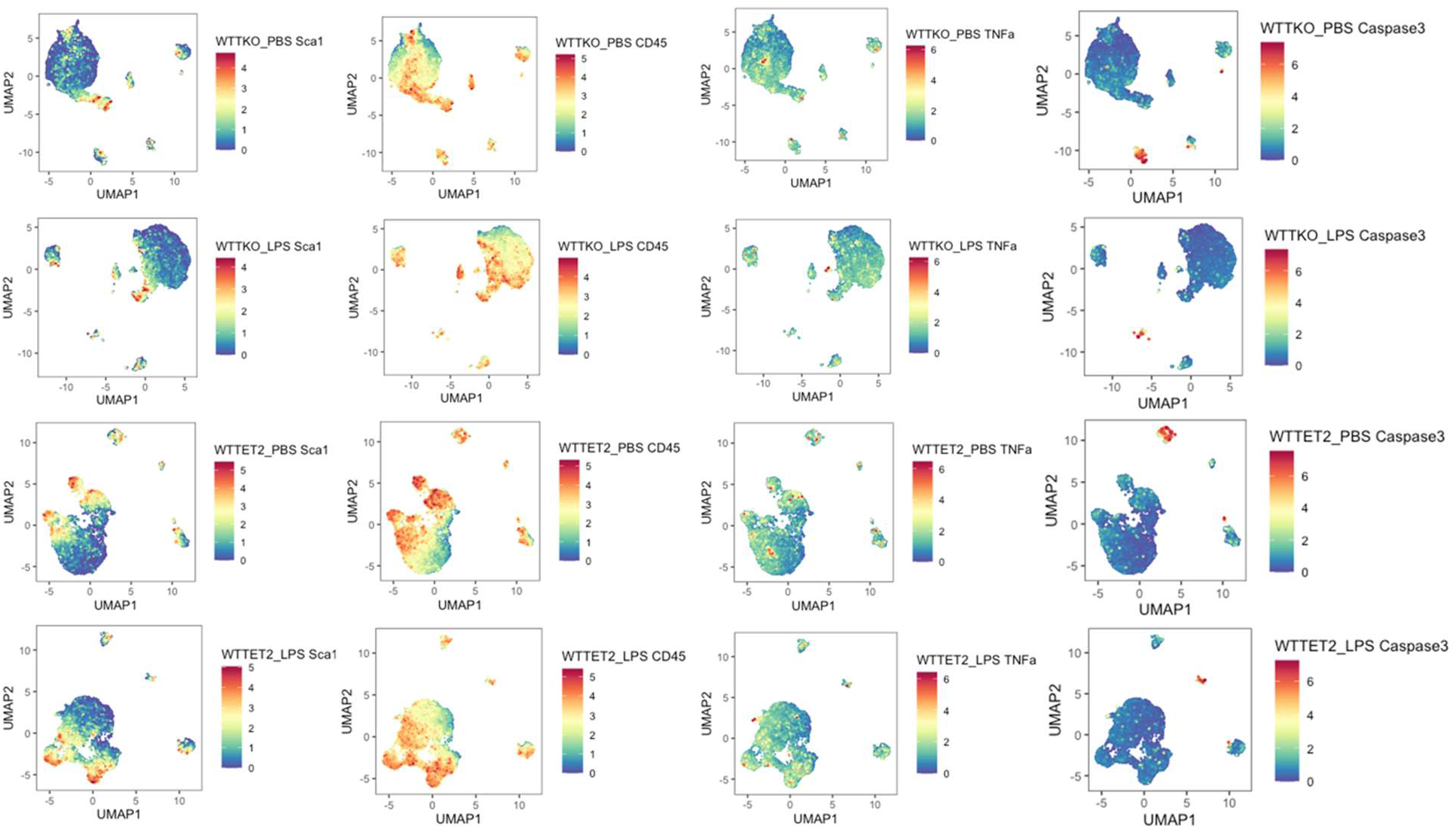
Differential cell count illustrated from the unsupervised learning. Markers of interest under genetic perturbation, including Sca1 (LSK cells), CD45 (leukocytes), TNFα (cytokines), and Caspase 3 (cell death indicator), were mapped to the clustering results from the T-REX algorithm.

**Table S1.** Cell markers investigated in this study

| Marker Type | Cell Type | Markers |
| --- | --- | --- |
| Lineage | Stem Cells | CD41, CD48, CD150, cKit, Sca1 |
| Lineage | Pan Leukocytes | CD45 |
| Lineage | T Lymphocytes | CD3c, CD8a, CD4, CD25 |
| Lineage | Mature Myeloid Cells | CD11b, Gr1 |
| Lineage | B Lymphocytes | CD43, IgM, CD45R |
| Lineage | Erythrocytes | Ter119 |
| Functional | Cytokines | TNFα, IL2, IL6 |
| Functional | Cell Death | Caspase3, Granzyme B |
| Functional | Proliferation | Ki67 |
| Pre-gating |  | DNA1, DNA2 |
| Instrument |  | Cell Length, Time |

**Table S2.** Searching for the optimal size “hyperparameter” via RNN-LSTM model architecture

| Micro Sample Size | Train Acc | Val Acc | Test Acc |
| --- | --- | --- | --- |
| 2 | 0.73653 | 0.71342 | 0.71033 |
| 10 | 0.93403 | 0.88667 | 0.87375 |
| 30 | 0.99791 | 0.95369 | 0.95125 |
| 60 | **0.99499** | **0.9325** | **0.9575** |
| 120 | 1.0 | 0.9598 | 0.975 |
| 240 | 1.0 | 0.93 | 0.93 |
| 480 | 1.0 | 0.77551 | 0.88 |

**Table S3.**
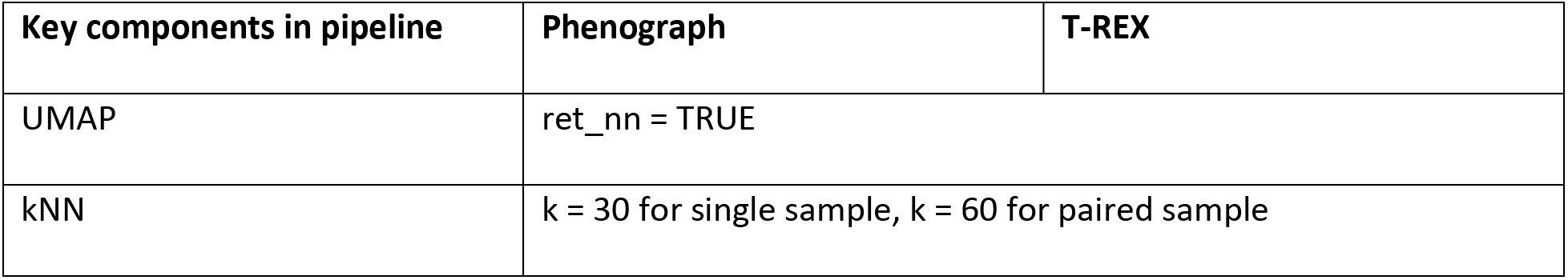
Hyperparameters applied in PhenoGraph and T-REX

## Notes

### Competing Interest Statement

The authors have declared no competing interest.

### Summary of Updates

Main text updated to clarify results; Introduction revised to improve clarity; Figure legends revised.

